# Quantitative method for estimating stain density in electron microscopy of conventionally prepared biological specimens

**DOI:** 10.1101/2019.12.11.873323

**Authors:** Andrea Fera, Qianping He, Guofeng Zhang, Richard D. Leapman

## Abstract

Stain density is an important parameter for optimizing the quality of ultrastructural data obtained from several types of 3D electron microscopy techniques, including serial block-face electron microscopy (SBEM), and focused ion beam scanning electron microscopy (FIB-SEM). Here, we show how some straightforward measurements in the TEM can be used to determine the stain density based on a simple expression that we derive. Numbers of stain atoms per unit volume are determined from the measured ratio of the bright-field intensities from regions of the specimen that contain both pure embedding material and the embedded biological structures of interest. The determination only requires knowledge of the section thickness, which can either be estimated from the microtome setting, or from low-dose electron tomography, and the elastic scattering cross section for the heavy atoms used to stain the specimen. The method is tested on specimens of embedded blood platelets, brain tissue, and liver tissue.

## Introduction

A new generation of imaging techniques in the electron microscope provide biologists with three-dimensional nanoscale ultrastructure that helps to elucidate mechanisms for a wide range of important cellular processes. Two such techniques are serial block-face electron microscopy (SBEM) (Denk & Horstmann, 2004; Briggman et al., 2011; Helmstaedter et al., 2013; Pfeifer et al., 2015; Shomorony et al., 2015; Porovskaya et al., 2018, Porovskaya et al., 2019), and focused ion beam scanning electron microscopy (FIB-SEM) (Hekking et al., 2009; Drobne, 2013; Narayan & Subramaniam, 2015; Glancy et al., 2015). In both these approaches, embedded blocks of cells or tissues, are imaged in a scanning electron microscope using the backscattered electron signal generated by scattering of electrons from heavy-atom stain incorporated into the biological specimens after fixation and prior to embedding. In the SBEM a diamond knife built into the specimen stage successively cuts thin layers from the block and the block face is imaged after each layer is removed, whereas in the FIB-SEM an ion beam removes thin layers from the block face. FIB-SEM provides higher resolution perpendicular to the block face, but data acquisition is much faster in the SBEM, which enables the collection of 3–D images from volumes as large as 10^6^ µm^3^. The pixel size in the plane of the block face is ∼5 nm in the SBEM, but the minimum cutting increment, i.e., resolution perpendicular to the block face, is ∼25 nm, or more typically ∼50 nm, which produces highly anisotropic voxels. The FIB-SEM can provide nearly isotropic voxels of size ∼5nm, but from smaller volumes of cells and tissues.

Unlike specimens prepared for thin section TEM, it is not possible to post-stain the block, so that all the heavy-metal stain must be incorporated into the block before data are acquired. Optimization of sample preparation is crucial for obtaining the best results from SBEM and FIB-SEM (Deerinck et al., 2010). Too little stain produces images with poor signal-to-noise ratio, which reduces visibility of ultrastructure, yet too much stain can mask subtle ultrastructural features. In the acquisition of 3D images of ultrastructure using SBEM, the fluence of incident electrons is limited to approximately 20 electrons/nm^2^ due to shrinkage of the block caused by radiation damage. If higher doses are used, the specimen block will not cut evenly at smaller cutting increments (Kizilyaprak et al., 2015).

Insufficient concentration of stain in specimens prepared for SBEM can lead to electrical charging of the block, which can severely distort the images stacks and make the data unusable. It is possible to mitigate charging effects to some extent by filling the SEM specimen chamber with a partial pressure of gas, which ionizes under electron irradiation, and thereby provides a source of ions that can neutralize charge build-up in the specimen. However, introduction of gas into the SEM column reduces the backscattered image quality, due to electron scattering by the gas molecules. Recently, a focal charge compensation device has been developed that delivers a much lower concentration of gas through a thin tube positioned just above the specimen block (Deerinck et al., 2018).

In a previous study (Sousa et al., 2008), it was shown that by collecting annular dark-field images in the scanning transmission electron microscope (STEM) at two different collection angles it is possible to determine quantitatively the distributions of heavy-atoms of stain and light-atoms of embedding resin and biological structures, based on differences in the angular distributions of elastically scattered electrons by the light and heavy atoms. In those experiments, the different collections angles were obtained by varying the camera length so that the annulus of the detector subtended different scattering angles. Measurements made on an osmium-fixed, but otherwise unstained, freeze-substituted preparation of the unicellular marine green alga, *Ostreococcus tauri*, showed that the sample contained 1.2 ± 0.1 osmium atoms nm^−3^. However, that approach depended on setting up STEM imaging for different well-defined camera lengths and making careful measurements from thin specimens. Here, we demonstrate a more straightforward technique for determining stain density in embedded biological specimens using conventional bright-field TEM imaging of sections that are cut from the same blocks that are subsequently analyzed by SBEM or FIB-SEM. The robustness of the method is demonstrated for sections cut at a thickness from 100 nm to 750 nm, and the same stain density is obtained regardless of the specimen thickness, or whether there is mass loss caused by electron irradiation, and the measurements and analysis can be performed in a few minutes.

## Materials and Methods

### Specimen Preparation

Mouse liver tissues were gifts from Drs. Rodney L. Levine and Lo Lai, National Heart, Lung, and Blood Institute (NHLBI), and from Dr. Bechara Kachar, National Institute on Deafness and Other Communication Disorders (NIDCD), NIH. Procedures for obtaining the tissue were approved by the NHLBI Animal Care and Use Committee (Protocol #H0120) and performed in accordance with the guidelines described in the Animal Care and Welfare Act (7 USC 2144). Specimen preparation. Sample blocks of mouse liver were prepared for SBEM using the University of California San Diego (UCSD) NCMIR protocol. Pieces of liver were dissected from mice and were fixed in a mixture of 2.5% glutaraldehyde and 2% formaldehyde in sodium cacodylate buffer with 2 mM calcium chloride at room temperature for 5 minutes, and for an additional 2–3 hours on ice in the same solution. The specimens were washed three times with cold cacodylate buffer containing 2 mM calcium chloride and post-fixed for 1 hour in a solution of reduced osmium containing 2% osmium tetroxide, 1.5% potassium ferrocyanide 2 mM CaCl_2_ in 0.15 mM sodium cacodylate buffer (pH 7.4), followed by incubation with a 1% thiocarbohydrazide (TCH) solution in ddH_2_O for 20 minutes at room temperature. Subsequently, samples were fixed with 2% OsO_4_ in ddH_2_O for 30 minutes at room temperature, followed by 1% aqueous uranyl acetate at 4 °C for 12 hours. The samples were then subjected to *en bloc* Walton’s lead aspartate staining (Walton, 1979), and placed in a 60°C oven for 30 minutes. Following dehydration in graded concentrations of ice-cold anhydrous ethanol for 5 minutes each, the specimens were placed in anhydrous ice-cold acetone at room temperature for 10 minutes. Tissue pieces were infiltrated with 30% Hard-Plus resin for 2 hours, 50% Hard-Plus for a further 2 hours, 75% Hard-Plus overnight, and 100% Hard-Plus for one day before being polymerized in a 60°C oven for 48 hours.

Epon-embedded blocks of mouse brain tissue were prepared using a similar protocol that is described above for the liver specimen with initial fixation in paraformaldehyde and glutaraldehyde and stained with heavy metals (OsO_4_ used as a fixative, uranyl acetate, and lead aspartate) (Deerinck et al., 2010).

Blocks of human blood platelets in the resting state were prepared for thick-section scanning TEM (STEM) tomography as described previously (Pokrovskaya et al., 2015; McBride et al., 2018). In brief, blood was drawn using institutional review board–approved procedures (Sehgal and Storrie, 2007). Anticoagulant-containing blood was fixed with 3% paraformaldehyde and 0.1% glutaraldehyde immediately after draw using a stock solution prepared in phosphate-buffered saline. Following isolation, platelet preparations were fixed a second time in 2.5% glutaraldehyde, washed with 0.1 mol/L sodium cacodylate buffer for 5 min on ice and post-fixed with 0.8% K_3_Fe(CN)_6_, 1.0% OsO4 for 30 min at room temperature. After further washes, the samples were stained with 0.5% uranyl acetate for 1h at room temperature, and were embedded in Epon-12 epoxy resin. This sample preparation procedure resulted in blocks that were lightly stained and suitable for STEM tomography. In the present study, we were interested in comparing the stain concentration with sample blocks prepared with the UCSD NCMIR protocol used to acquire 3-D data in the SBEM (Deerinck et al., 2010).

Normally, these blocks would be imaged in the SBEM for serial block-face scanning electron microscopy, but for the present purposes, the blocks were mounted in a Leica EM UC6 Ultramicrotome, and sections of known but varying thickness (100 nm, 250 nm, 500 nm, and 750 nm) were cut and deposited on 3-mm copper mesh electron microscopy grids.

### TEM measurements

Specimens were imaged with an FEI/Thermo Fisher Scientific Tecnai TF30 TEM, equipped with a twin objective lens and a field-emission source, using a Gatan Ultrascan 4000 CCD camera. This instrument was operated at a beam energy of 300 keV because we wanted to test our method on samples up to 1 µm in thickness, which would not have been possible with a TEM operating at a beam energy of 100 keV or 120 keV. However, the measurements we have described could be obtained from thinner specimens using any TEM equipped with a CCD camera.

Regions of embedded liver, brain, and blood platelets were selected, which contained clear regions of embedding resin without the presence of stained cells or tissues. The sections were imaged at low magnification to obtain fields approximately 10 µm across, which contained stained cells as well as clear plastic. An objective aperture of diameter 40 µm was selected, which accepted scattering angles up to *β* = 10 mrad, as expected from an objective of radius *r*_*oa*_ = 20 µm, and focal length *f*_*obj*_ = 2 mm in the Tecnai TF30 TEM, which gives a collection angle *r*_*oa*_ / *f*_*obj*_ = 10 mrad. The size of the collection angle *β* was confirmed by recording the diffraction pattern from a polycrystalline gold film, with known diffraction rings, with and without the objective aperture inserted.

### Determination of stain concentration

The transmitted intensity through a plastic section of thickness *t* containing embedded heavy-atom stained cells is given by:

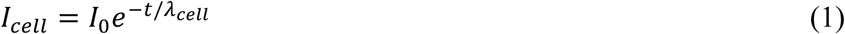

where *I*_0_ is the incident electron fluence, *I*_*cell*_ is the transmitted fluence through the cell, and *λ*_*cell*_ is the elastic mean free path for the specific region of interest in the embedded cell.

Rewriting Eq. (1) gives:

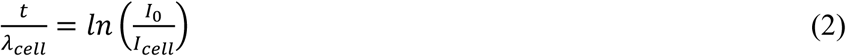

Similarly, for regions of pure resin in a nearby regions with the same thickness:

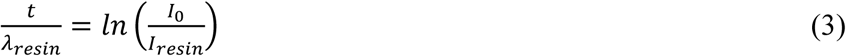

where *λ*_*resin*_ is the elastic mean free path for the pure resin.

We consider that the stained biological structures are embedded in a block of epoxy resin containing *n*_*i*_ atoms of type *i* per unit volume, and the elastic scattering cross section of atom *i* is *σ*_*i*_ The elastic mean free path for the pure epoxy resin is then given by:

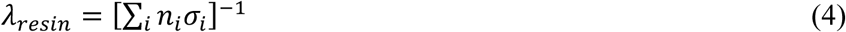

Further, we consider that the biological structures are stained with heavy stain atoms of type *h* containing *n*_*h*_ per unit volume distributed sparsely throughout the embedded cells or tissues, and the elastic scattering cross section of heavy atom *h* is *σ*_*h*_. The elastic mean free path for the embedded cell is then given by:

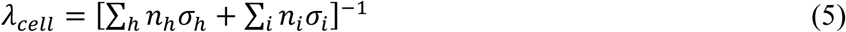

Based on the observed lack of contrast in unstained embedded biological structures, we assume here that the unstained biological material does not attenuate the primary electron beam due to elastic scattering more than the surrounding clear (unstained) resin. This is expected since the light atoms in the resin have approximately the same mass as the atoms in the biological tissue or cells. However, more precision could be achievable by accounting for the difference in elastic scattering between the resin and the biological material.

Substituting Eq. (2) and Eq. (3) into Eq. (4) and Eq. (5), we obtain:

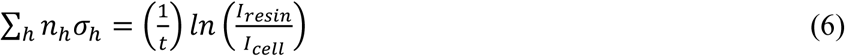

Specimens prepared for 3D SBEM and FIB-SEM are typically stained with osmium (atomic number 76), lead (atomic number 82), and uranium (atomic number 92). Previously, we have acquired energy-dispersive x-ray spectra from specimens prepared for SBEM and observed all three elements in the stained blocks, but we found that lead had the highest concentration (He et al., 2018).

If the total number of heavy atoms *H* per unit volume is *n_H_* = Σ*_h_n_h_*, and the fraction of heavy atoms of each type *h* is *f_h_* = *n_h_*/*n*_H_, we can write Eq. (6) as

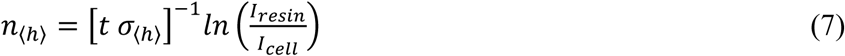

In general, the fraction of heavy atoms *f_h_* of type *h* is not known exactly and might vary somewhat from region to region in the specimen. To estimate the uncertainty in the number per unit volume of heavy atoms *n_H_* of all types in Eq. (7), we determine the variation in elastic scattering cross section as a function of atomic number for the different types of heavy atoms that are incorporated into the specimen (i.e., Os, Pb and U).

Since the regions of pure resin and regions of resin that contain stained biological material suffer the same mass loss under electron irradiation, Eq. (7) is almost independent of radiation exposure. The only requirements are (1) knowledge of the specimen thickness, which can be determined from the microtome settings for the sectioning, or from low dose electron tomography of sections on which nanoparticles have been deposited as fiducial markers, and (2) knowledge of the elastic scattering cross sections, which are now known with higher precision than was possible in earlier work, as shown below.

### Elastic scattering cross sections

Earlier work on computation of elastic scattering cross sections (Reimer & Ross-Messemer, 1989) shows the dependence on atomic number *Z*, with *σ*_*el*_ is proportional to *Z*^*4/3*^ in the Lenz model (Lenz, 1954), and proportional to *Z*^*3/2*^ in the Thomas-Fermi model (Schafer et al., 1971; Langmore, 1973). Recently, more accurate determinations have been made that provide differential scattering cross sections as a function of scattering angle *θ* (Salvat et al., 2005), and these computations are now available as a Standard Reference Database SRD 64 from the National Institute of Standards and Technology (Salvat et al., 2002). We make use of these data in the present study.

The differential elastic cross sections as a function of scattering angle *θ* for incident electrons with an energy of 300 keV are shown in Fig. 1 for atoms of (A) osmium, (B) lead, and (C) uranium. These plots were obtained by converting solid-angle differential cross sections to angle-differential cross sections using the data provided in the NIST Standard Reference Database SRD 64 (Salvat et al, 2002; Jablonski et al., 2004; Salvat et al., 2005):

**Figure 1.**
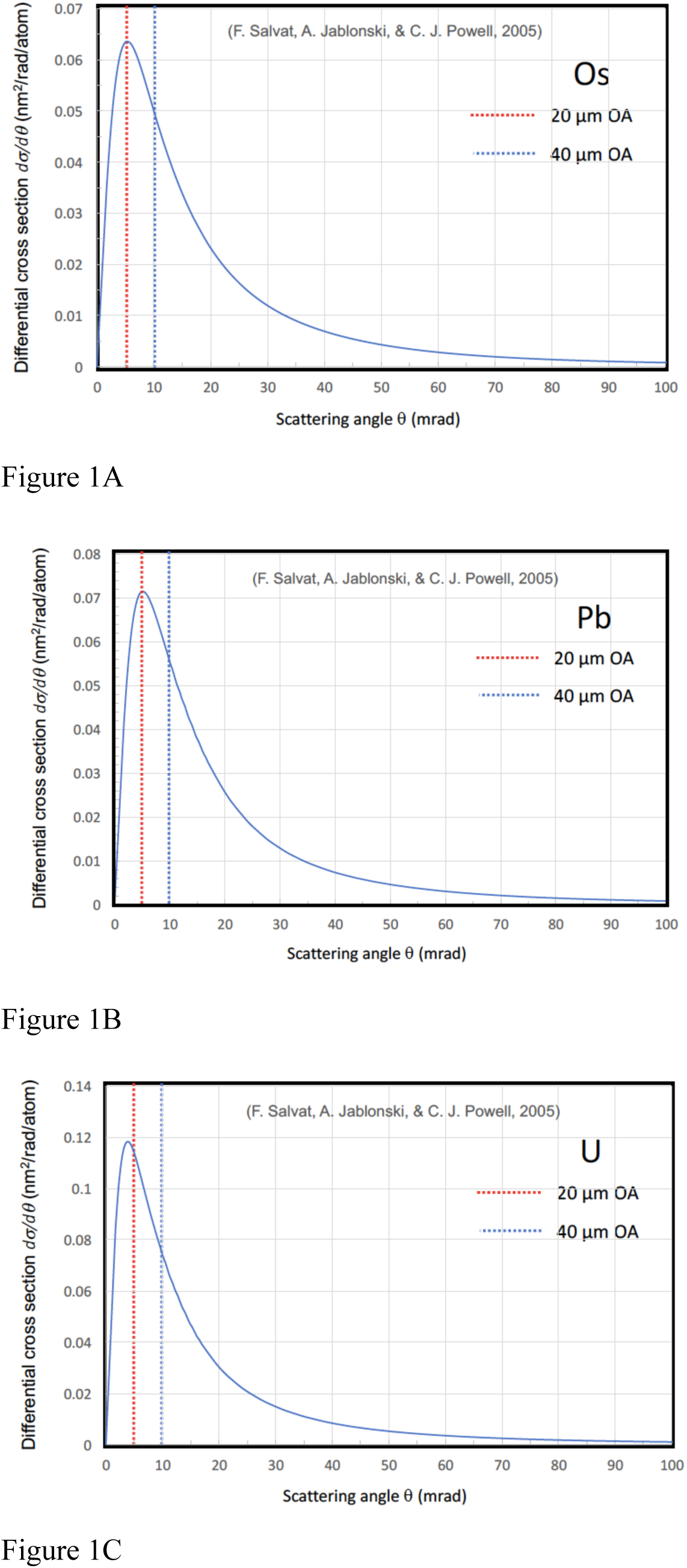
Differential elastic scattering cross sections, as a function of polar scattering angle, for incident electrons of energy 300 keV for (A) osmium, (B) lead, and (C) uranium atoms. This is taken from the NIST standard reference database SRD 64 (ref), and as described by Salvat et al., 2005. Angles for two objective apertures in the FEI/Thermo Fisher Scientific Tecnai TF30 TEM are indicated: a 20-µm diameter aperture subtending scattering angles of 0.005 radian, and a 40-µm diameter aperture subtending scattering angles of 0.01 radian.

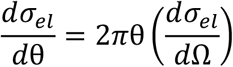

where Ω is the solid angle, θ is the polar angle, *d*Ω is a small element of solid angle, and *d*θ is a small element of polar angle.

The partial cross section for scattering outside of the objective aperture subtending semi-angle β is given by integrating the polar-angle differential cross section over polar angles from β to π.

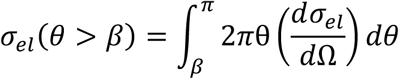

The cross sections for elastic scattering of 300-keV incident electrons by osmium, lead, and uranium atoms are presented in Table 1 for β → 0, i.e., total cross section, and for two objective aperture sizes in the Tecnai TF30 TEM: β = 5 mrad (20 µm diameter), and β = 10 mrad (40 µm diameter), i.e., partial cross sections. We selected the 40-µm aperture to minimize possible contributions of multiple inelastic scattering in very thick specimens (>500 nm). Values for the total elastic cross sections are consistent with the values from the earlier work, but the model used to compute the cross sections for the NIST SRD 64 database is expected to be more accurate since it is based on a relativistic (Dirac) partial-wave calculations for scattering by a local central interaction potential.

**Table 1.** Elastic cross sections for 300-keV electrons scattering elastically from osmium, lead and uranium atoms: total cross section (column 2); and partial elastic scattering cross sections for objective aperture diameters of 20 µm equivalent to a semi-angle β = 0.005 radian (column 3), and 40 µm equivalent to a semi-angle β = 0.01 radian (column 4), which is selected for the present study.

| Element | $\sigma_{\text{el}}(\beta \rightarrow 0)$<br>($\text{nm}^2$) | $\sigma_{\text{el}}(\beta = 0.005)$<br>($\text{nm}^2$) | $\sigma_{\text{el}}(\beta = 0.01)$<br>($\text{nm}^2$) |
| --- | --- | --- | --- |
| Os | 0.00123 | 0.00103 | 0.00077 |
| Pb | 0.00137 | 0.00114 | 0.00085 |
| U | 0.00186 | 0.00144 | 0.00102 |

## Results and discussion

Bright-field images containing 2048 × 2048 pixels were acquired with a Gatan Ultrascan CCD camera on a FEI/Thermo Fisher Scientific Tecnai TF30 TEM at a beam energy of 300 keV and with a 40-µm diameter objective aperture passing scattering angles *β* = 0.01 radians. The raw bright-field images are presented in the panels on the left sides of Fig. 2 for the human platelet sample, Fig. 3 for the mouse brain sample, and Fig. 4 for the mouse liver sample; the intensities of the images are represented as *I*(*x, y*), where *x, y* are the coordinates of the pixels, and the intensities are in CCD counts. The panels on the right side of each figure are corresponding computed values of *ln*[*I*_*resin*_/*I*(*x, y*)], where *I*_*resin*_ is the mean intensity of a region of clear resin contained in the image. Although the raw images are in arbitrary units, the pixel values of the *ln*[*I*_*resin*_/*I*(*x, y*)] image are not in arbitrary units because the logarithm is independent of the scaling of the *I*(*x, y*) image.

**Figure 2.**
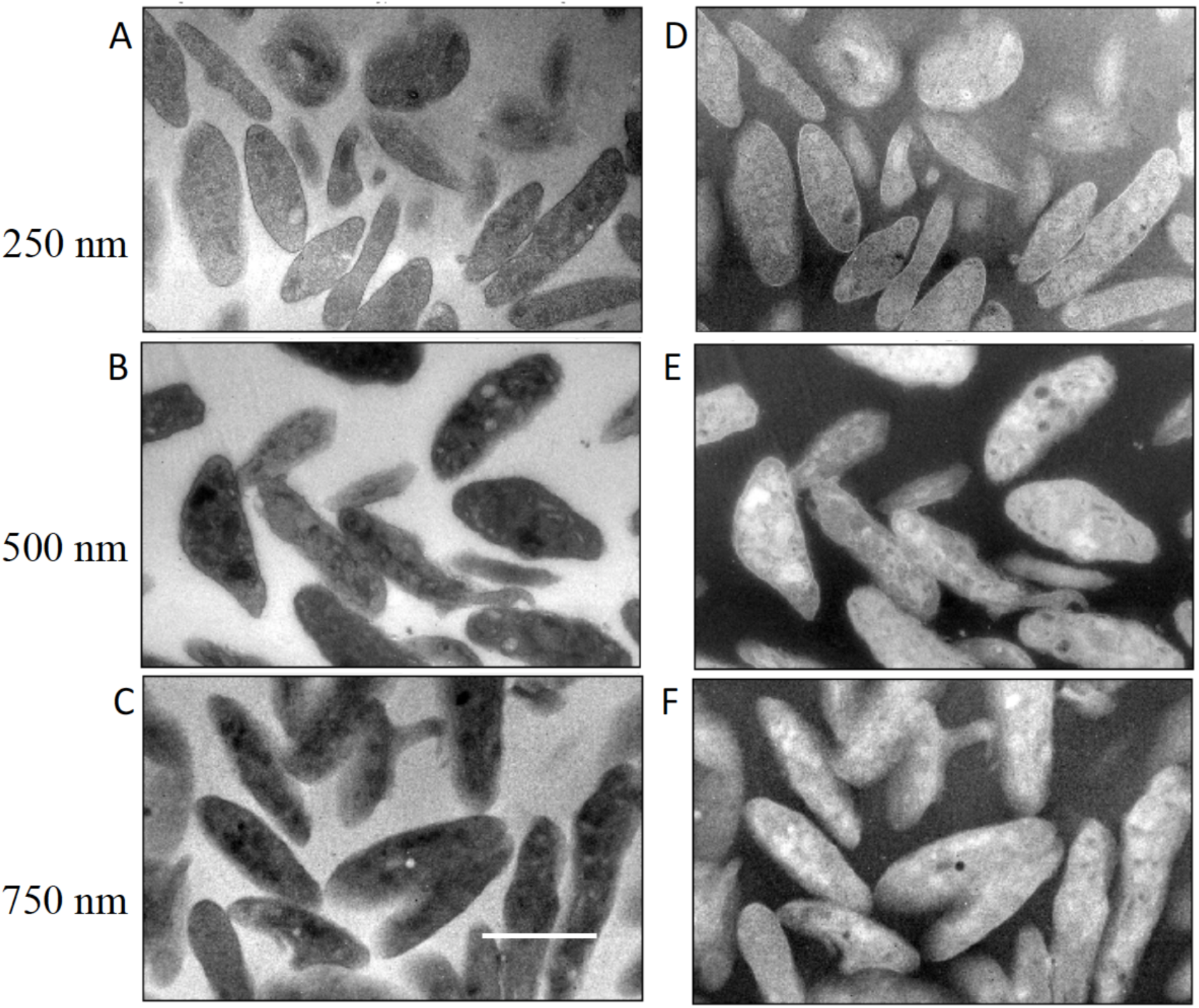
Analysis of bright-field TEM images *I*(*x, y*) from sections of human blood platelets in specimens of different thicknesses: (A) 250 nm; (B) 500 nm; (C) 750 nm. Corresponding computed images of *ln*[*I*_*resin*_/*I*(*x, y*)] are shown in (D) for the 250-nm section; (E) for the 500-nm section; and (F) for the 750-nm section. Values for *I*_*resin*_ were obtained by averaging the intensity of the pixels contained in regions of clear embedding resin surrounding the cells. Scale bar = 2 µm.

**Figure 3.**
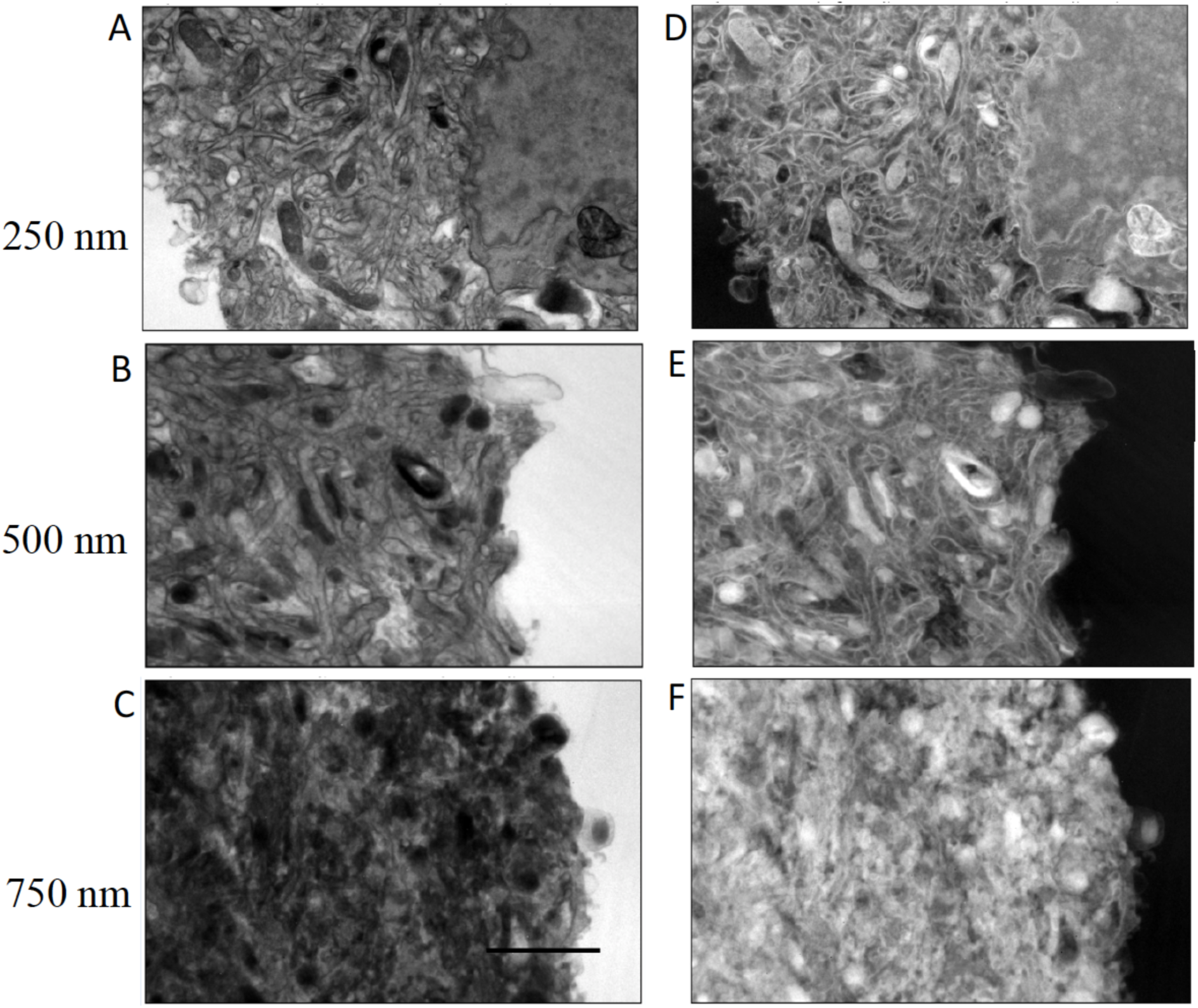
Analysis of bright-field TEM images *I*(*x, y*) from sections of mouse brain in specimens of different thicknesses: (A) 250 nm; (B) 500 nm; (C) 750 nm. Corresponding computed images of *ln*[*I*_*resin*_/*I*(*x, y*)] are shown in (D) for the 250-nm section; (E) for the 500-nm section; and (F) for the 750-nm section. Values for *I*_*resin*_ were obtained by averaging the intensity of the pixels contained in regions of clear embedding resin surrounding the cells. Scale bar = 2 µm.

**Figure 4.**
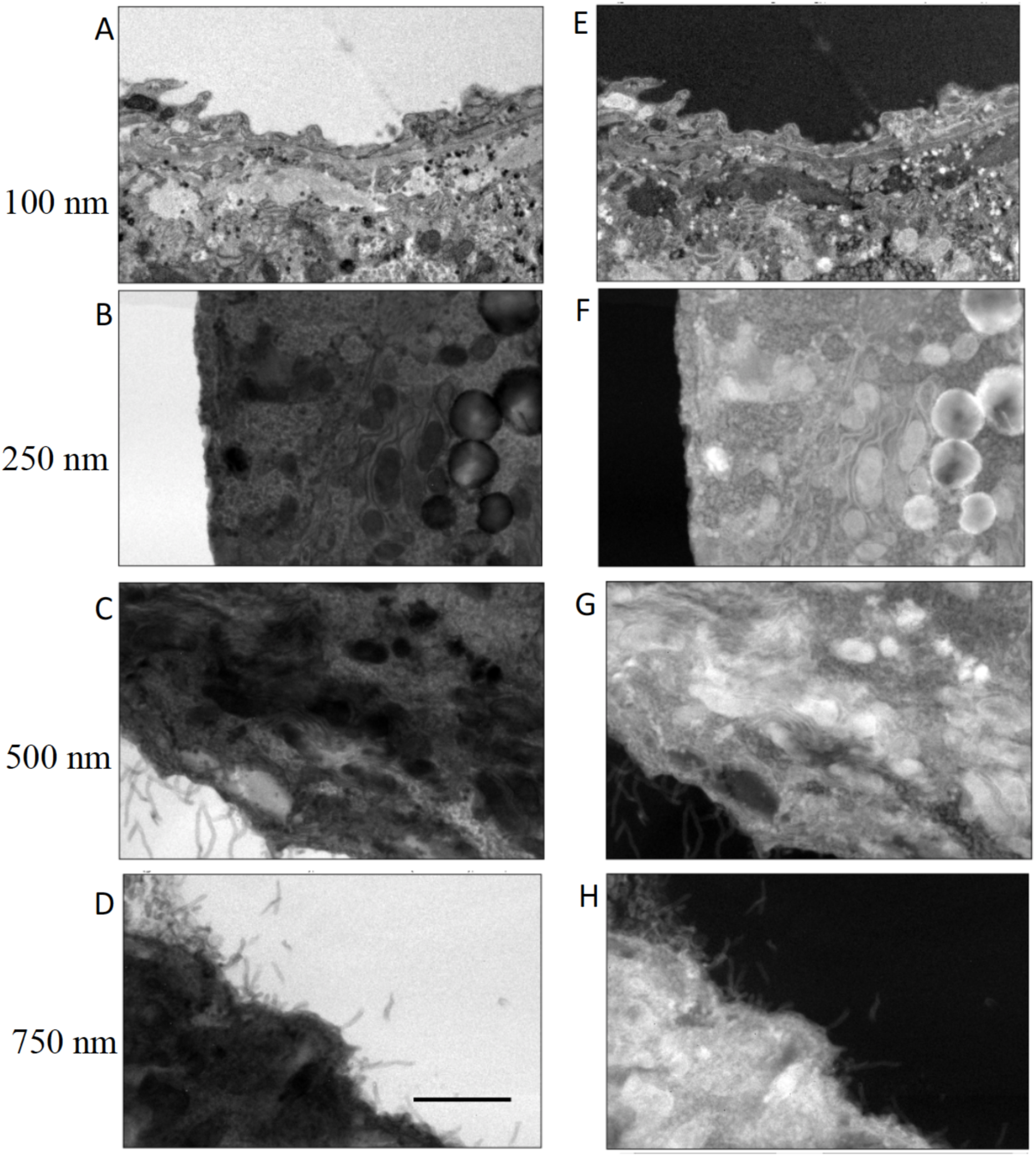
Analysis of bright-field TEM images *I*(*x, y*) from sections of mouse liver of different thicknesses: (A) 100 nm; (B) 250 nm; (C) 500 nm; (D) 750 nm. Computed images of *ln*[*I*_*resin*_/*I*(*x, y*)] are shown in (E) for a 100-nm section; (F) for a 250-nm section; (G) for a 500-nm section; and (H) for a 750-nm section. Values for *I*_*resin*_ were obtained by averaging the intensity of pixels contained in regions of clear embedding resin surrounding the cells. Scale bar = 2 µm.

Multiple measurements of ⟨*ln*[*I*_*resin*_/*I*_*cell*_(*x, y*)]⟩ were made from mean values of *ln*[*I*_*resin*_/ *I*(*x, y*)] in different specimen regions for samples of thickness 250 nm, 500 nm, and 750 nm. For the liver specimen (Fig. 4), measurements were also performed for a section of 100-nm thickness.

Plots of ⟨*ln*[*I*_*resin*_/*I*_*cell*_(*x, y*)]⟩ versus section thickness are presented in Fig. 5A for the human platelet sample, in Fig. 5B for the mouse brain sample, and in Fig. 5C for the mouse liver sample. It is evident that the plots for the platelet sample and the mouse brain sample show a linear relation with a straight line passing through the origin. There is some evidence that the liver sample departs from linearity for the thickest (750-nm) specimen. This departure is not surprising since the liver sample contains the highest stain concentration, and in view of the thickness of the section.

**Figure 5.**
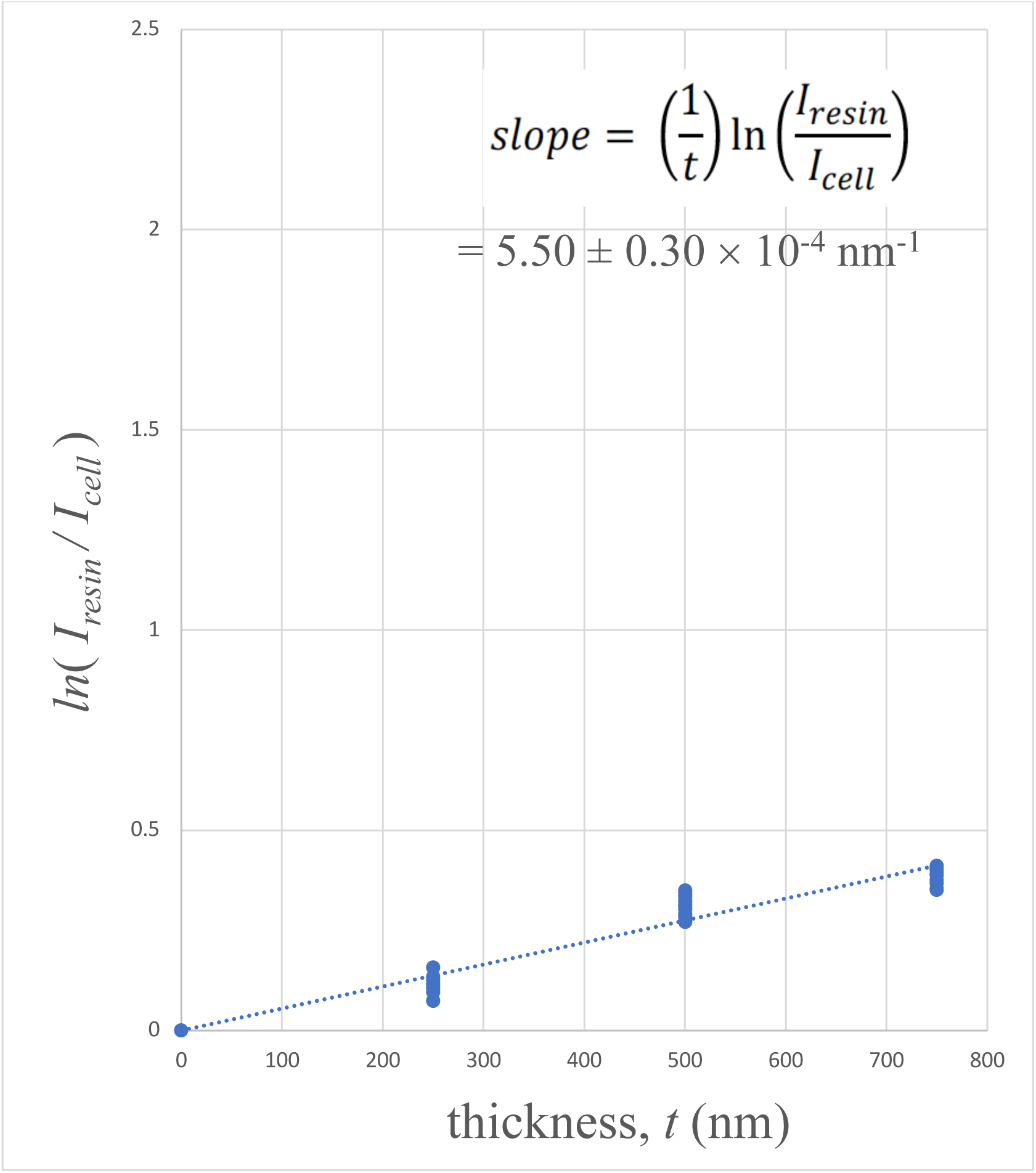
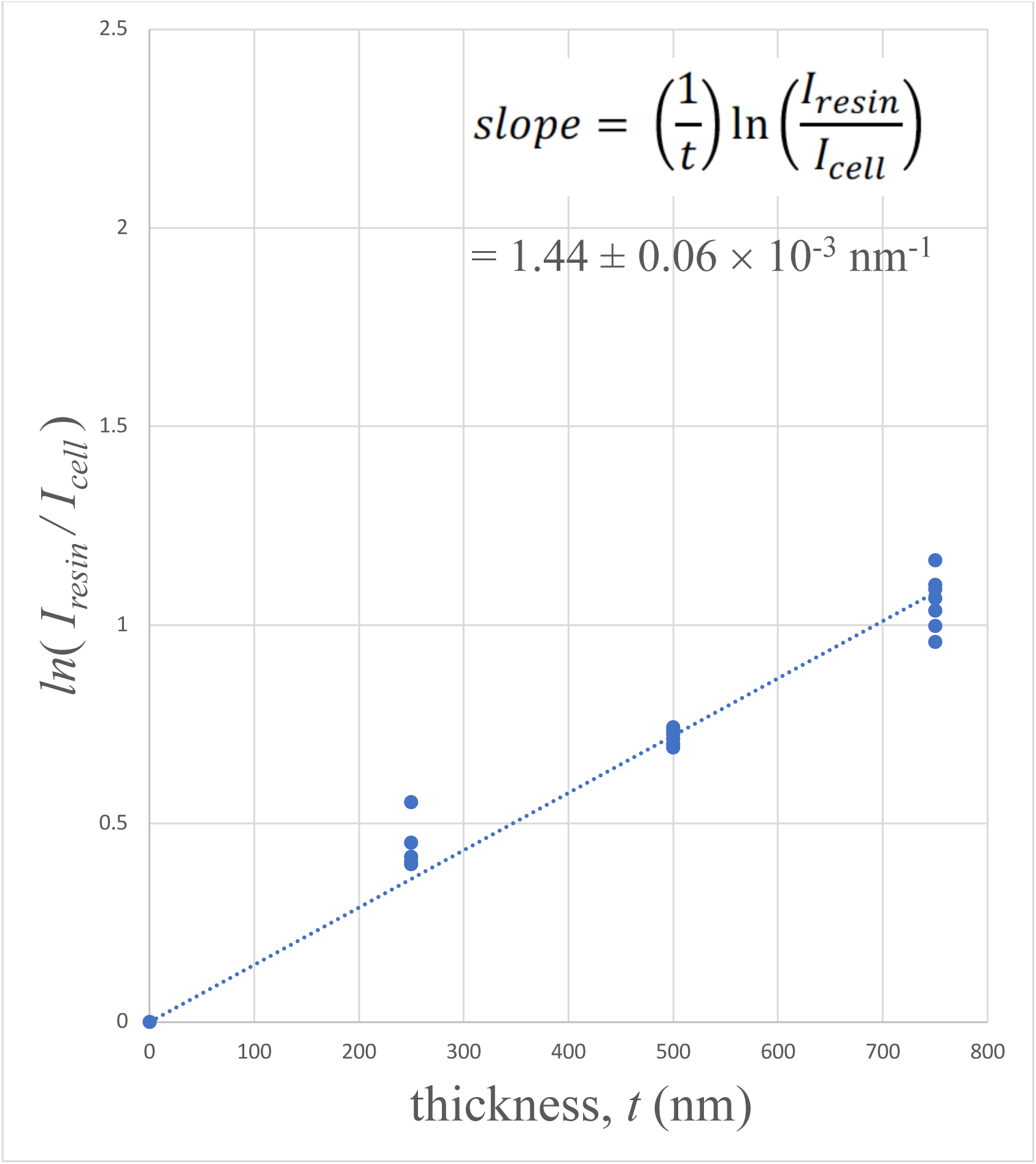
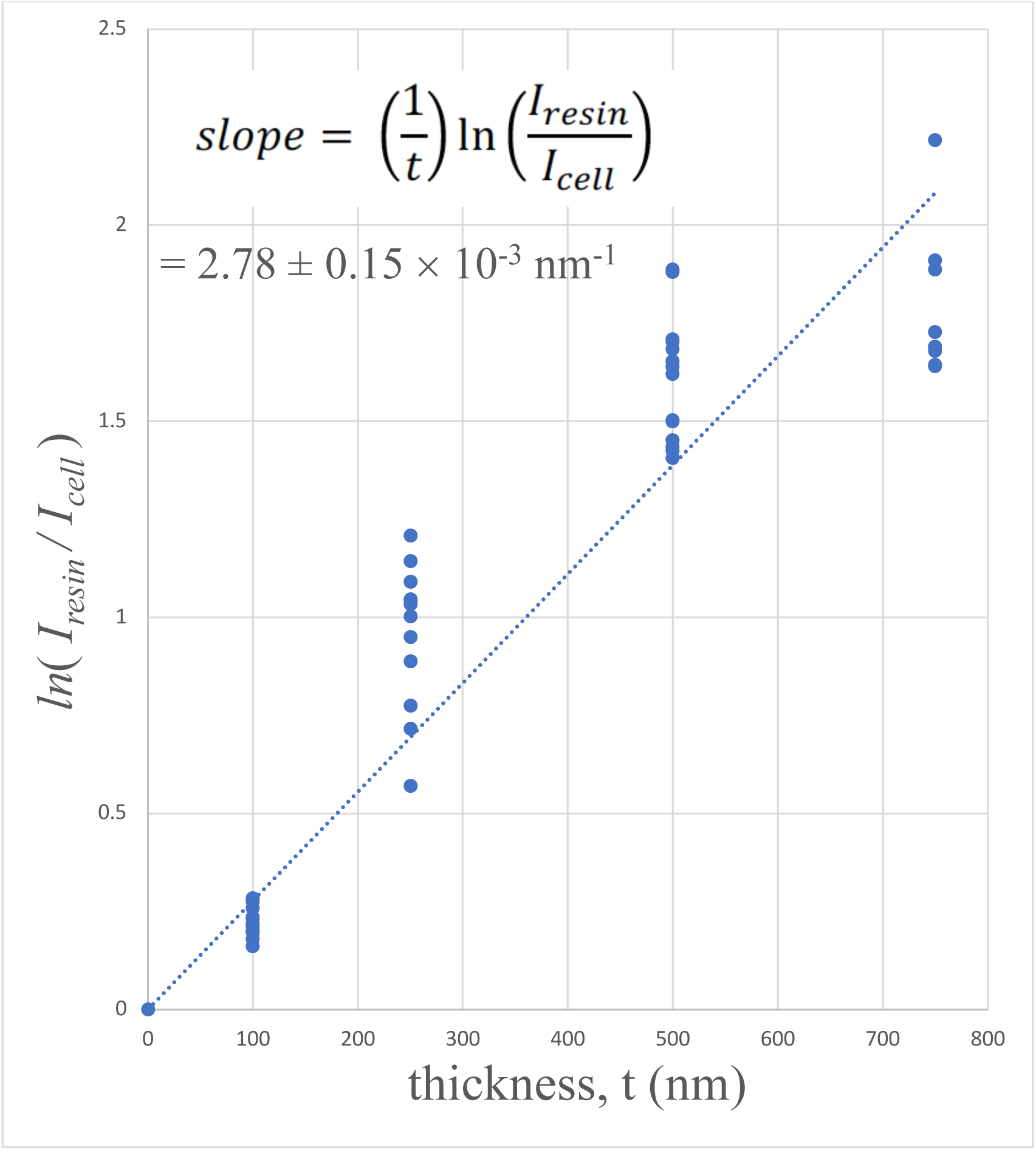
Plots of ⟨*ln*[*I*_*resin*_/*I*_*cell*_(*x, y*)]⟩ versus specimen thickness *t* for: (A) human blood platelet sample; (B) mouse brain sample; and (C) mouse liver sample. The linear fits were constrained to pass through the origin since very thin specimens must contain negligible numbers of heavy stain atoms.

The stain concentration can be determined by dividing the slope of each plot by the average elastic scattering cross section for the heavy atoms present in the specimen. When information is known about the stain atoms, e.g., if only Os or only U is used to stain a specimen, the appropriate elastic scattering cross section in Table 1 can be used. Although the human blood platelet sample was stained with a low concentration of uranyl acetate as well as OsO_4_, we have assumed that osmium was the dominant heavy element in this specimen, and we have therefore taken the experimentally determined slope of the plot in Figs. 5A, together with the Os partial elastic scattering cross section, 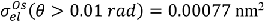 to estimate a stain density of 0.71 ± 0.04 Os atoms nm^−3^ (Table 2, row 1). The concentration of heavy atoms in the lightly stained sample of blood platelets is commensurate with the measurements of osmium stain concentration in the freeze-substituted preparation of *Ostreococcus tauri* obtained by STEM imaging by Sousa et al. (2008).

**Table 2.**
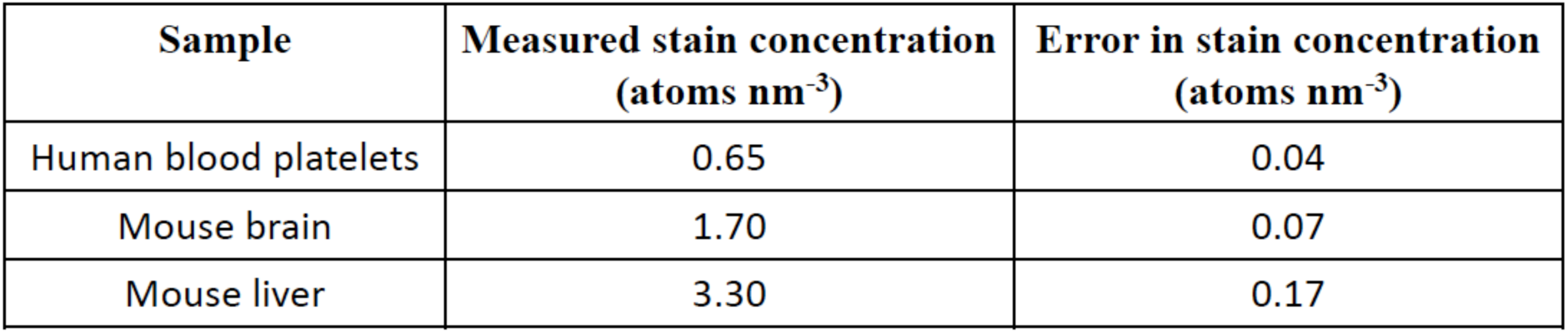
Measured stain concentration in heavy atoms per nm^3^ for sections of human blood platelets, mouse brain, and mouse liver. The error estimates are based on the standard deviation in the measured slopes of the plots of ⟨*ln*[*I*_*resin*_/*I*_*cell*_(*x, y*)]⟩ versus thickness. The uncertainty in the elastic scattering cross section is not considered, since the values are expected more accurate than the variations in the slopes.

We have previously used energy-dispersive x-ray spectroscopy to measure relative concentrations of Os, Pb and U in SBEM blocks prepared with the UCSD NCMIR protocol (He et al, 2018). In those specimens, we found that Pb is the most abundant heavy atom. Since the atomic number of Pb lies between that of Os and U in the periodic table, and Pb is also the most abundant element, we take the Pb partial elastic scattering cross section, 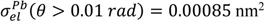 (Table 1), to estimate the stain concentration. Table 1 shows that the elastic cross section for Os is ~9% less than that for Pb, and the elastic cross section for U is ~20% higher than that for Pb, which is the major element in the mouse brain and mouse liver specimens. Using the partial elastic scattering cross section for Pb, together with experimentally determined slopes of plots in Figs. 5B and 5C, we obtain a stain density of 1.70 ± 0.07 heavy atoms nm_-3_ for the mouse brain sample, and 3.3 ± 0.17 heavy atoms nm_-3_ for the mouse liver sample (Table 2, rows 2 and 3).

Since the plots in Figs. 5A-C were obtained from independent measurements from specimens cut to different thicknesses, the uncertainties in the specimen thickness are accounted for by the calculated errors in the slopes of *ln*(*I_resin_*/*I_cell_*) vs. *t*, if there are no systematic errors in the section thickness due to miscalibration of the ultramicrotome or compression of the sections.

The density of epoxy resin is 1.22 gm/cm^3^, and if it is assumed that the mean atomic mass of the non-hydrogen atoms in the resin is the same as for carbon, i.e., 12 g/mole, the epoxy resin contains approximately 60 non-hydrogen atoms per nm^3^. The blood platelet sample therefore contains a heavy-to-light atom stain concentration of 1.0 atomic %, a relatively low value that is expected since this specimen was not prepared for SBEM analysis. This level of staining would likely be suboptimal for SBEM analysis and could result in electrical charging, unless gas were introduced into the specimen chamber of the SEM, or unless a focal charge compensation device were employed. Even if electrical charging were prevented, the backscattered electron images would be noisy.

The specimen of mouse brain contains a heavy-to-light atom concentration of 2.8 atomic %, which is typical for specimens prepared with the UCSD NCMIR protocol.

The specimen of mouse liver contains a heavy-to-light atom concentration of 5.5%, which is higher than for most specimens prepared with the UCSD NCMIR protocol. It is likely that such high stain concentrations could result in loss of ultrastructural information

## Conclusion

We have derived a simple expression for determining the concentration of heavy atoms of stain from bright-field TEM images of resin embedded biological specimens that are prepared for 3-D imaging by SBEM or FIB-SEM. The method could also be used to determine stain concentrations in any other electron microscope imaging mode, and it depends only on knowledge of the specimen thickness and the elastic scattering cross sections for stain atoms (most commonly, osmium, lead and uranium). The specimen thickness can be obtained from the setting of the ultramicrotome, or if this estimate is not sufficiently precise, the thickness can be determined from low-dose tomography tilt series from specimens on which gold nanoparticles are deposited on both surfaces. We have made use of cross sections, available from the NIST standard reference database SRD 64 (Salvat et al., 2005), which are based on relativistic (Dirac) partial-wave calculations, and which can be computed for any elements of interest and for any beam energy, and objective aperture size.

We have tested the method on three specimens with different levels of staining and the results appear to be consistent; blocks prepared with the UCSD NCMIR protocols have the highest stain concentration, and samples prepared for thick-section STEM tomography have a stain concentration that is a factor of between 3–5 times smaller.

Due to logarithmic term in Eq. (6), the number of light atoms from the resin and biological material do not appear in the expression for the stain concentration, and the number of heavy atoms per unit volume is computed directly. The inverse product of the thickness and the scattering cross section in Eq. (6) has units of nm^−3^, so that the measurement needed to determine the stain concentration is a dimensionless quantity that is independent of the light atoms. This makes the method very robust with respect to mass loss during bright-field imaging, or contamination deposited on the section while being imaged in the column of the TEM.

In our experience, considerable time is wasted by imaging specimens in which the stain concentration is not well suited to the imaging technique, and it is often not possible to assess the stain concentration by inspection of the specimen in the optical microscope, since even at low concentration osmium causes black staining of the block, whereas lead and uranium are not easily visible. The method described here can be applied very quickly, simply by recording a few bright field images. Moreover, it is not necessary to use a 300-keV TEM, and measurements can be made just as easily at beam energies of 100 or 120 keV.

## Acknowledgments

This work was supported by the Intramural Research Program of the National Institute of Biomedical Imaging and Bioengineering. The authors thank Dr. Brian Storrie (University of Arkansas for Medical Sciences, Little Rock, AR, USA) for the blood platelet specimen, and Dr. Rodney L. Levine and Dr. Lo Lai, National Heart, Lung, and Blood Institute, NIH, for the gift of the mouse liver sample.

